# A comparison of methodological approaches to the study of young sex chromosomes: A case study in *Poecilia*

**DOI:** 10.1101/2021.11.29.470452

**Authors:** Iulia Darolti, Pedro Almeida, Alison E. Wright, Judith E. Mank

## Abstract

Studies of sex chromosome systems at early stages of divergence are key to understanding the initial process and underlying causes of recombination suppression. However, identifying signatures of divergence in homomorphic sex chromosomes can be challenging due to high levels of sequence similarity between the X and the Y. Variations in methodological precision and underlying data can make all the difference between detecting subtle divergence patterns or missing them entirely. Recent efforts to test for X-Y sequence differentiation in the guppy have led to contradictory results. Here we apply different analytical methodologies to the same dataset to test for the accuracy of different approaches in identifying patterns of sex chromosome divergence in the guppy. Our comparative analysis reveals that the most substantial source of variation in the results of the different analyses lies in the reference genome used. Analyses using custom-made *de novo* genome assemblies for the focal species successfully recover a signal of divergence across different methodological approaches. By contrast, using the distantly related *Xiphophorus* reference genome results in variable patterns, due to both sequence evolution and structural variations on the sex chromosomes between the guppy and *Xiphophorus*. Changes in mapping and filtering parameters can additionally introduce noise and obscure the signal. Our results illustrate how analytical differences can alter perceived results and we highlight best practices for the study of nascent sex chromosomes.

## Introduction

Substantial recent attention in sex chromosome research has focused on the earliest stages of X-Y divergence in order to glean the initial processes of recombination suppression (Wright et al. 2016). Studies of nascent sex chromosome divergence will by definition result in subtle patterns of X-Y sequence differentiation as substantial differences have not yet sufficiently accumulated. Given the expected subtlety, methodology and underlying data can be quite important, and small changes may make all the difference between identifying a delicate pattern or missing it entirely.

For example, several recent tests for divergence between the guppy X and Y chromosomes have revealed contradictory results. Full genomic analysis of the *Poecilia reticulata* sex chromosomes was originally presented in Wright et al. (2017) based on comparisons between male and female genomes (Fig. 1). This approach can be used to identify what, if any, regions of the Y chromosome are diverged from the X, and to compare across populations to determine intra-specific variation. Wright et al. (2017) found a relatively small region (10Mb) of significant Y degeneration, designated Stratum I. This region was characterized by a reduction in the number of male reads that mapped compared to females, consistent with the concept of Y degeneration. Moreover, the same pattern was observed in all six of the natural populations assayed as well as a captive lab population, and the rules of parsimony therefore suggest that Stratum I is ancestral to the colonization of Trinidad. Wright et al. (2017) also observed evidence of a second region of nascent divergence, Stratum II, that appeared to have formed independently in three upstream populations, but was smaller in downstream populations. This region was characterized by an increase in male single nucleotide polymorphism (SNP) density compared to females but no degradation of the Y. This pattern is consistent with either greatly reduced or complete loss of male recombination in this region, or selection against recombinant males.

**Figure 1.**
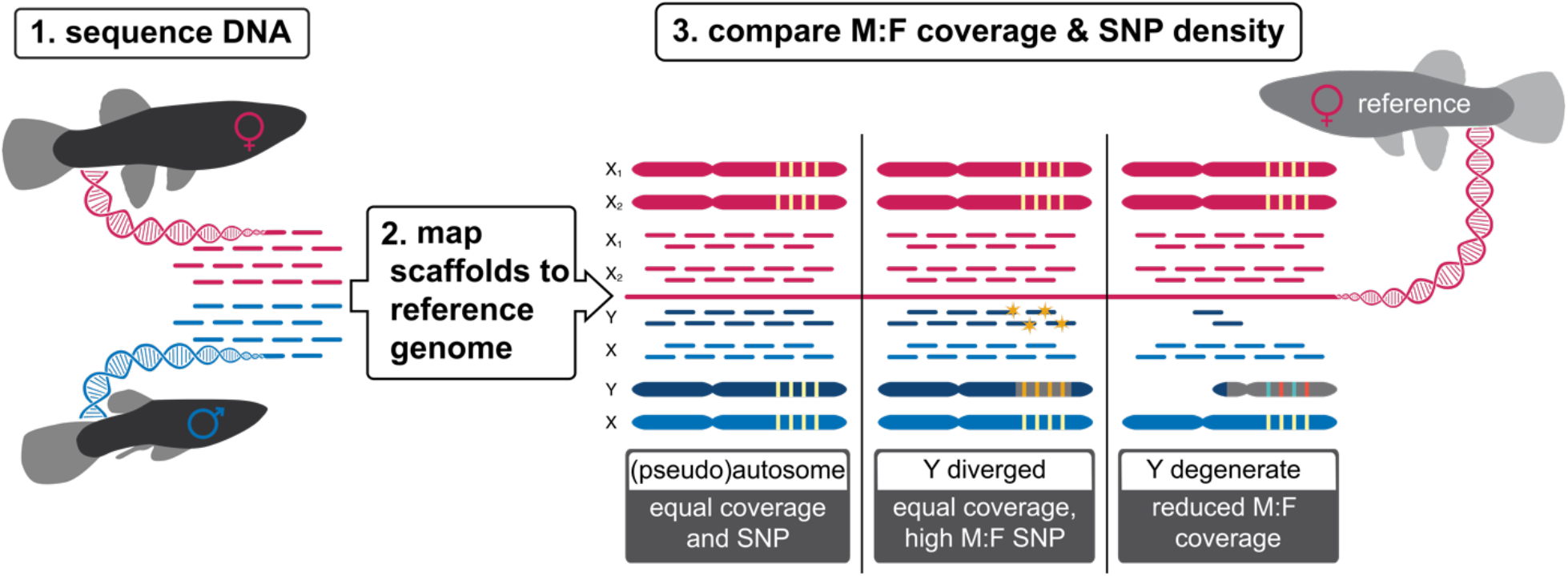
Genomic comparisons of male and female DNA data can be used to identify X-Y divergence. Step (1) Multiple males and females are sequenced, and female reads (red) are assembled, with resulting scaffolds ordered and oriented to the nearest available full reference genome. Step (2) Male (blue) and female (red) reads are mapped to this assembly. Step (3) Y divergence leads to male-specific SNPs, and therefore elevated male:female SNP density. As the Y degenerates, Y reads will no longer map to the X chromosome assembly, leading to reduced male:female coverage. Method adapted from Vicoso & Bachtrog 2013 and Vicoso and Bachtrog 2015. Figure courtesy of Jacelyn Shu (jacelyndesigns.com).

Almeida et al. (2021) built on these initial findings with a greatly expanded dataset, again recovering concordant patterns of Stratum I across the same six natural populations of *P. reticulata*. The expanded dataset incorporated 10X genomics linked-reads, allowing for far more sophisticated analyses. Namely by phasing X and Y haplotypes, it was possible to discern that Stratum I is comprised of two smaller separate regions of reduced male:female read depth. This region is also enriched for male-specific sequences, male-specific SNPs, and repetitive elements, the presence of which necessitate recombination suppression from the X chromosome. Importantly, there was also evidence of phylogenetic clustering of phased Y sequence in this region, indicating ancestral recombination suppression. Finally, this replication recovered evidence of parallel expansion of Stratum II in upstream populations.

Expanding phylogenetically, Darolti et al (2019) uncovered consistent patterns of sex chromosome topology in *P. wingei*. Initial karyotype analysis suggested that the X and Y chromosomes are ancestral to the common guppy (*P. reticulata*) and Endler’s Guppy (*P. wingei*) (Nanda et al. 1993). Furthermore, Darolti et al. (2019) found the same small region of Y chromosome degeneration consistent with Stratum I, although somewhat more pronounced in the degree of divergence from the X than *P. reticulata*. The region of degeneration matched nearly perfectly with *P. reticulata*, suggesting Stratum I was in fact present in the common ancestor of *P. wingei* and *P. reticulata*. Consistent with this, Morris et al (2018) found evidence of male-specific sequence shared between *P. reticulata* and *P. wingei*, possible only if recombination between the X and Y was halted in the common ancestor of these species. Moreover, Darolti et al. (2020) used SNP segregation patterns from RNA-seq data across pedigrees to determine X and Y sequence, and found four genes that showed phylogenetic evidence of recombination suppression in the ancestor of *P. wingei* and *P. reticulata*. Although the bootstrap values for any one locus were not excessively high, it is telling that all four were in Stratum I. The ancestral origin of Stratum I was further supported by conserved patterns of male-hypomethylation within this region in both species (Metzger et al. 2020), consistent with sexualization of gene regulation. Finally, Darolti et al. (2019) found evidence for another independent origin of Stratum II based on SNP data in *P. wingei*. Work in outgroup species revealed the same chromosome is a sex chromosome in *P. picta* and *P. parae* (Darolti et al. 2019; Sandkam et al. 2021), although diverged to a far greater degree in both these species.

Crucially, all of these analyses were based on custom genome or transcriptome assemblies generated bespoke from the underlying data (Wright et al. 2017; Darolti et al. 2019; Darolti et al. 2020; Almeida et al. 2021), although they did use existing related reference genomes to physically place and orient scaffolds. This is in contrast to other studies which have used existing resources derived from different populations or species, resulting in potential mismatches between the underlying data and the genome to which it is compared. Taking a bespoke approach is critical as it reduces the phylogenetic distance between the sequence reads and the reference to which they are mapped, which can increase the proportion of reads that are accurately mapped and reduce issues arising from structural variation and repetitive sequence. Secondly, an important step in identifying diverged regions in sex chromosomes is ensuring stringent mapping parameters (Caravalho and Clark 2013; Smeds et al. 2015; Vicoso and Bachtrog 2013; Vicoso and Bachtrog 2015; Palmer et al. 2019). This is particularly relevant for homomorphic sex chromosomes as they still retain sequence orthology between the X and Y, and incorrectly mapped reads can mask coverage differences between the sexes and lead to the misclassification of sex-linked sequences as autosomal. Wright et al. (2017), Darolti et al. (2019) and Almeida et al. (2021) used stringent mapping limits, removed minor alleles with low frequency, which likely represent sequencing errors, and focused on coding sequence to minimize issues with repetitive elements (Table 1). This was based on the reasoning that young sex chromosomes would exhibit subtle divergence signatures, and stringency would be required to detect it (Palmer et al. 2019; Vicoso and Bachtrog 2015).

**Table 1.** A comparison of methods and findings for *P. reticulata* sex chromosome strata.

| Study | Evidence for Stratum I from M:F read depth | Phylogenetic clustering of Y sequences in Stratum I | Evidence for Stratum II from M:F $F_{ST}$ | Genome Assembly | Read depth analysis | SNP analysis |
| --- | --- | --- | --- | --- | --- | --- |
| Wright et al. (2017) | Yes | n/a | Yes | Bespoke (*P. reticulata*) | Uniquely mapping reads | Limited to coding sequence; Site coverage > 10; SNP frequency > 0.3 x site coverage |
| Bergero et al. (2019) | No | n/a | Yes | Kunstner et al. (2016) (*P. reticulata*) | Default mismatch parameters, duplicate reads excluded | Quality score > 30; Minimum coverage 20; Biallelic SNPs |
| Darolti et al. (2019) | Yes | n/a | Yes | Bespoke (*P. reticulata* and *P. wingei*) | Uniquely mapping read | Limited to coding sequence; Site coverage > 10; SNP frequency > 0.3 x site coverage |
| Charlesworth et al. (2020) | No | n/a | Yes | Kunstner et al. (2016) (*P. reticulata*) | Not reported* | Not reported* |
| Fraser et al. (2020) | Yes | n/a | No | Bespoke (*P. reticulata*) | Default mismatch parameters | MAF > 0.05 |
| Kirkpatrick et al. (2020) | No | Yes | No | Schartl et al. (2013) (*Xiphophorus maculatus*) | Default read mapping parameters with local argument, duplicate reads excluded | Quality score > 20; Minimum read depth 3; Biallelic SNPs |
| Almeida et al. (2021) | Yes | Yes | Yes | Bespoke (*P. reticulata*, river-specific) | Uniquely mapping reads, duplicate reads excluded | MAF > 0.1; excluding extremely high coverage sites |

\*Mapping criteria not reported in methods and bioinformatic code not publicly available. Defaults assumed. MAF = minor allele frequency.

Despite observing remarkable concordance of these patterns across multiple datasets, species and analytical methods, other recent studies have differed substantially in their approach and reported some different results (Table 1). For example, Bergero et al. (2019) did not report evidence for Stratum I in their own *P. reticulata* data, and although they did uncover a pattern that is broadly consistent with Stratum II, it was not statistically different across populations (Charlesworth et al. 2020). Fraser et al. (2020) identified small male-specific regions largely consistent with the regions identified by Almeida et al. (2021), although because of scaffold orientation differences and population-specific inversions, they are in different physical locations. Finally, Kirkpatrick et al. (2020), reanalyzing data from Darolti et al. (2019) found evidence of Stratum I and Stratum II in *P. wingei*, but not in *P. reticulata*. Notably, they did find phylogenetic evidence of recombination suppression in the ancestor of these two species in Stratum I, consistent with Darolti et al. (2020) and Almeida et al. (2021).

Importantly Bergero et al. (2019), Charlesworth et al. (2020) and Kirkpatrick et al. (2020) relied on existing reference genomes for all their analyses and did not use genomic reads to build custom-made assemblies for the target species. Importantly, Kirkpatrick et al. (2020) mapped reads from *P. wingei* and *P. reticulata* to *Xiphophorus*, which last shared a common ancestor 40 mya (Kumar et al. 2017). Additionally, Bergero et al. (2019) used less stringent mapping criteria, and Charlesworth et al. (2020) and Fraser et al. (2020) used default mapping parameters. Together, this produces two sources of potential methodological noise in replication efforts. First, noise can arise from accumulated mutations due to phylogenetic distance between the samples used to generate sequence reads and the genome that they are mapped to. Second, permissive default mapping parameters allow for mismapping, and therefore potentially result in significant noise in genomic comparisons between males and females. In addition to this, many of these studies used different underlying datasets that varied in sample origin, number and read depth, and so it is difficult to distinguish the role of sample variation from methodological differences in these discrepancies.

The proliferation of studies on this system with different levels of analytical sophistication allow for a remarkable comparison of the role of genomic methodology in pattern discovery. We tested various methods on the same underlying data with the goal of determining the methodological reasons for inconsistent findings across these studies, and to develop best practices moving forward to the genomic study of nascent sex chromosome systems.

## Methods

### Datasets

Using the *P. reticulata* data from Almeida et al. (2021) and the *P. wingei* data from Darolti et al. (2019), we ran multiple analyses of guppy sex chromosome evolution, following the various analytical methods used by Wright et al. (2017), Bergero et al. (2019) and Kirkpatrick et al. (2020), as summarized in Table 1 and detailed below. We were unable to include the methodology of Charlesworth et al. (2020) as mapping criteria was not reported in methods and bioinformatic code is not publicly available. The datasets for *P. reticulata* and *P. wingei* included paired-end DNA-seq reads from three males and three females from the Quare upstream population (EBI ENA under BioProject PRJEB39998) and from our lab population (NCBI SRA under BioProject PRJNA528814), respectively. We assessed read quality using FastQC v0.11.9 (www.bioinformatics.babraham.ac.uk/projects/fastqc/, last accessed 8 November 2021), trimmed using Trimmomatic v0.36 (Bolger et al. 2014) and concatenated reads as in Darolti et al. (2019) and Almeida et al. (2021). To replicate previous studies, all analyses were repeated using several different genomes and their respective gene annotations, which included the *P. reticulata* Quare *de novo* genome assembly from Almeida et al. (2021), the *P. reticulata* reference genome from Kunstner et al. (2016) (NCBI accession GCF_000633615.1), the *P. wingei de novo* genome assembly from Darolti et al. (2019) and the *Xiphophorus maculatus* reference genome from Schartl et al. (2013) (NCBI accession GCF_002775205.2, v5.0).

### Coverage analysis

For each focal species, we used three separate methodological pipelines to map and filter reads and to estimate read depth. The first method followed the analysis in Wright et al. (2017), which used bwa v0.7.15 aln/sampe (Li and Durbin 2009) to map reads, removed reads that were not uniquely mapping and estimated coverage with soap.coverage v2.7.7 (http://soap.genomics.org.cn, last accessed 1 April 2019). The second method followed the pipeline in Kirkpatrick et al. (2020), which mapped reads using bowtie2 v2.2.9 with default parameters and the -local argument (Langmead and Salzberg 2012), removed PCR duplicates using Picard v2.0.1 (http://broadinstitute.github.io/picard, last accessed 8 November 2021) and calculated coverage with BEDtools v2.26 (Quinlan and Hall 2010). Lastly, the third method followed the analysis in Bergero et al. (2019), which mapped reads with bwa mem and the -M argument (Li and Durbin 2009), removed PCR duplicates with BEDtools (Quinlan and Hall 2010) and estimated coverage using SAMtools v1.3.1 (Li et al. 2009).

For all three methodological pipelines, average coverage values were calculated separately for males and females, and average male:female coverage for each non-overlapping window was calculated as log_2_(average male coverage) – log_2_(average female coverage). A window size of 50kb was used for all *P. reticulata* analyses and *P. wingei* analyses based on the *X. maculatus* genome, while 10kb windows were used for *P. wingei* analyses using the more fragmented *de novo P. wingei* genome. Moving averages of coverage were plotted in R v4.0.5 (R Core Team 2019) based on sliding window analyses using the *roll_mean* function. Ninety-five percent confidence intervals for the moving average plots were obtained by randomly sampling autosomal values 1,000 times without replacement.

### SNP density analysis

To further assess patterns of Y divergence, for both *P. reticulata* and *P. wingei*, we compared three methodological approaches of estimating SNP density differences between males and females.

First, based on Wright et al. (2017), we mapped reads to each genome using bowtie2 with default parameters (Langmead and Salzberg 2012). After file sorting, we used bow2pro v0.1 (http://guanine.evolbio.mpg.de, last accessed 8 November 2021) to generate a profile for each sample, representing counts for each of the four nucleotide bases at each site. We then applied a minimum site coverage threshold of 10 and kept SNPs with a frequency of 0.3 times the site coverage. We further used gene annotation information to remove SNPs from the analysis if they were not located within coding sequences. For each sample, we calculated average SNP density for each gene as the sum of all SNPs divided by the sum of filtered sites in that gene, excluding those with zero filtered sites.

Second, following Kirkpatrick et al. (2020), we called variants from files previously filtered for PCR duplicates (see *Coverage analysis* section above) using BCFtools v.1.3.1 (Li 2011). We then filtered variants using VCFtools v0.1.12b (Danecek et al. 2011), removing indels and variants with a quality score lower than 20, and selecting for biallelic SNPs and a minimum read depth of 3. For each sample, we then used BEDtools counts to count the number of SNPs within 50kb windows across the genome.

Third, we used the pipeline in Bergero et al. (2019) to call SNPs from the PCR duplicates filtered files (see *Coverage analysis* section above) using GATK HaplotypeCaller v4.1.9 (Poplin et al. 2017) with the parameters --emit-ref-confidence GVCF and -stand-call-conf 30. Further genotyping was done with GATK GenotypeGVCFs with default parameters and SelectVariants to keep SNPs with a minimum coverage of 20, minimum quality of 30 and selecting for biallelic SNPs only. For each sample, we then used BEDtools counts to count the number of SNPs within 50kb windows across the genome.

Lastly, in each of these three methodological approaches, average SNP density across all males and across all females was calculated separately. For each gene or window, we calculated male:female SNP density as log_2_(average male SNP density) – log_2_(average female SNP density). We then divided male:female SNP density estimates into autosomal and sex-linked based on chromosomal position. The distributions of male:female SNP density for the autosomes and the sex chromosomes were plotted in R (R Core Team 2019) and differences between them were tested using Wilcoxon rank sum tests.

### Pairwise synteny analyses

We used LAST v1256 (Kielbasa et al. 2011) to perform pairwise synteny analyses between the *P. reticulata* sex chromosome (chromosome 12) from the reference genome (Kunstner et al. 2016), the *P. reticulata* sex chromosome from the Quare de novo assembly (Almeida et al. 2021) and the *X. maculatus* syntenic chromosome 8. For alignments involving the *X. maculatus* sequence, we used LAST with the HOXD70 seeding scheme designed for a higher rate of substitution, whereas for alignments involving *P. reticulata* sequences only we used the uNEAR seeding scheme for aligning sequences with lower rate of substitutions.

## Results

Using the same dataset across different genomes and methods, we first assessed the role of various genomic analysis parameters (Table 1) in detecting Stratum I on the *P. reticulata* and *P. wingei* sex chromosomes, previously reported in Wright et al. (2017), Darolti et al. (2019) and Almeida et al. (2021), summarized in Fig. 2 and Fig. 3.

**Figure 2.**
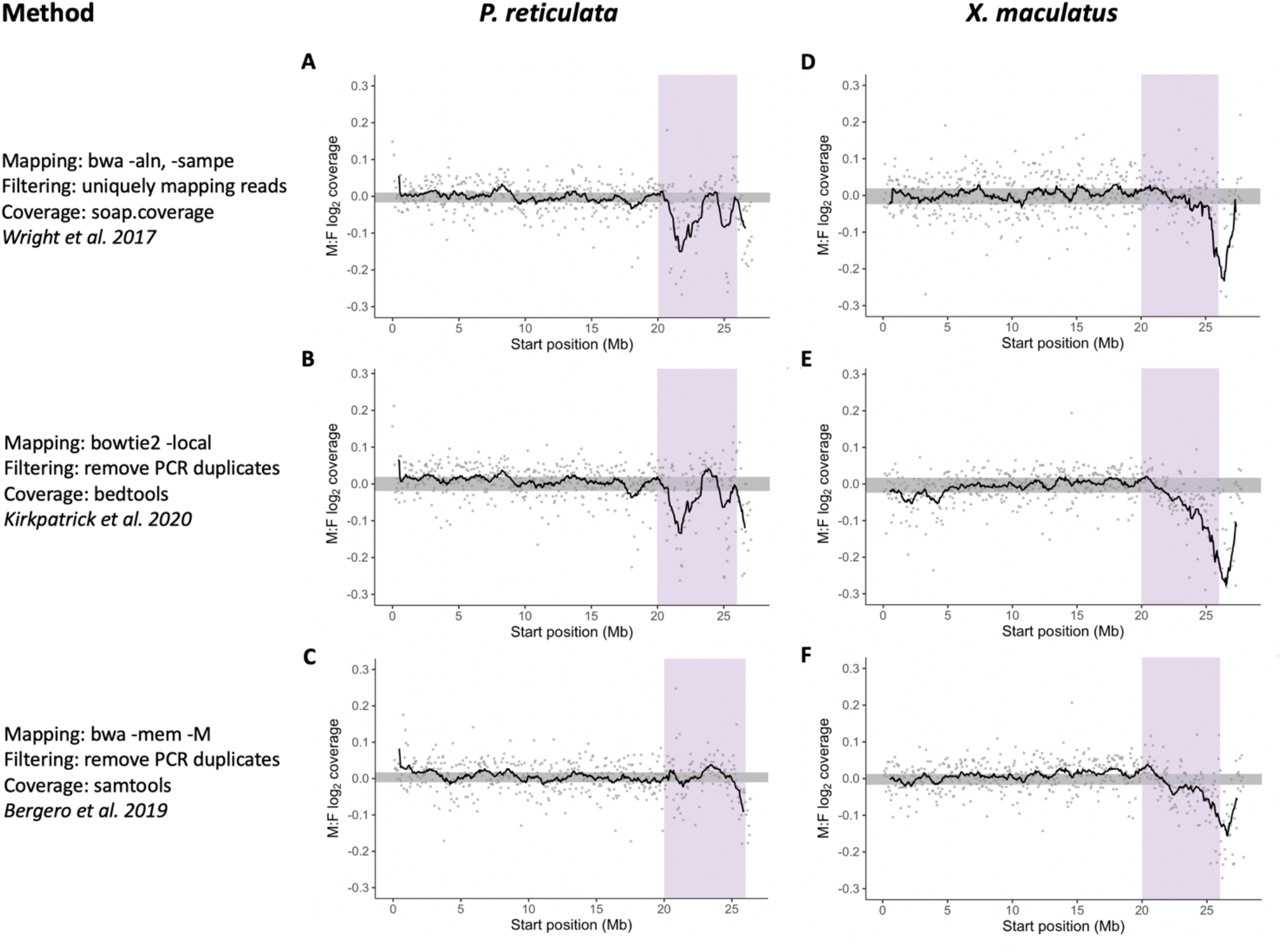
Signal for *P. reticulata* Stratum I using comparative methodological approaches. *P. reticulata* DNA-seq reads were mapped in turn to a *P. reticulata* genome assembly and the *X. maculatus* reference genome assembly (Schartl et al. 2013). For replicating previous studies, the *P. reticulata* reference genome from Kunstner et al. (2016) was used in the analysis based on the methods from Bergero et al. (2019), while the high quality Quare *de novo* assembly from Almeida et al. (2021) was used in the other two analyses. Moving average plots represent male to female coverage differences across the guppy sex chromosome (*P. reticulata* chromosome 12, and syntenic *X. maculatus* chromosome 8) in non-overlapping windows of 50kb. 95% confidence intervals, based on bootstrapping autosomal values, are shown in grey, and predicted boundaries for Stratum I from Almeida et al. (2021) are highlighted in purple.

**Figure 3.**
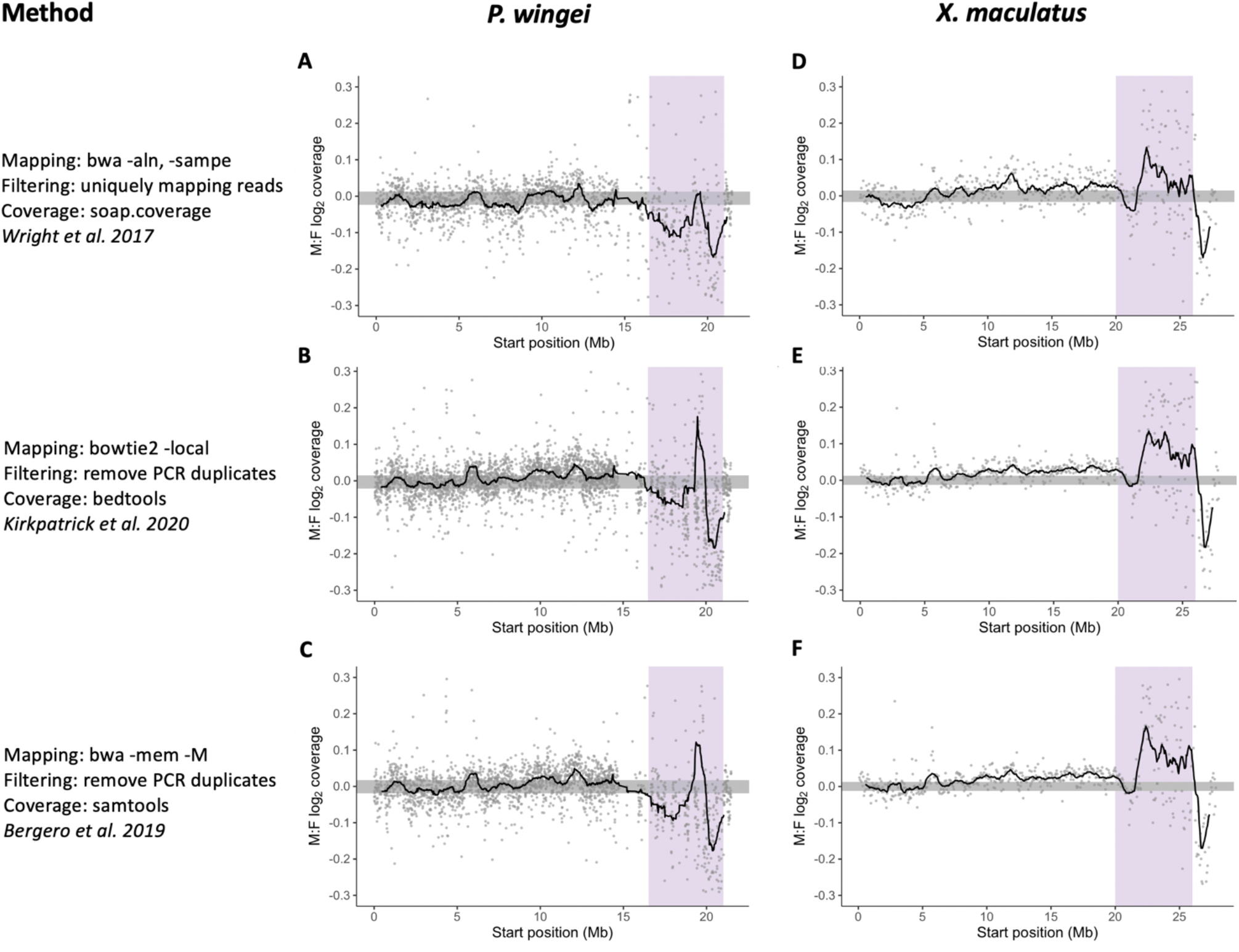
Signal for *P. wingei* Stratum I using comparative methodological approaches. *P. wingei* DNA-seq reads were mapped in turn to a *de novo P. wingei* genome assembly (Darolti et al. 2019) and the *X. maculatus* reference genome assembly (Schartl et al. 2013). Moving average plots represent male to female coverage differences across the sex chromosome (*P. wingei* chromosome 12, and syntenic *X. maculatus* chromosome 8) in non-overlapping windows of 50kb for the analyses that rely on the *X. maculatus* genome and windows of 10kb for the analyses that use the *de novo P. wingei* genome. The 95% confidence intervals, based on bootstrapping autosomal values, are shown in grey, and predicted boundaries for Stratum I from Darolti et al. (2019) are highlighted in purple.

Analyses on *P. reticulata* sequencing data that used the custom-made *de novo P. reticulata* genome assembly show a significantly lower male to female coverage, indicative of X-Y degeneration, at the distal end of the chromosome, in the previously estimated location of Stratum I (Fig. 2A, B). This pattern is evident from both the analysis that followed the pipeline from Wright et al. (2017) (Fig. 2A) and the analysis based on the methodology in Kirkpatrick et al. (2020) (Fig. 2B). All three analyses that relied on the *X. maculatus* reference genome also show a region with decreased male coverage relative to that in females, however, this region is shifted closer to the end of the chromosome and only partially overlaps with the syntenic region of the estimated location of *P. reticulata* Stratum I (Fig. 2D, E, F). Pairwise alignments revealed several structural rearrangements between the *P. reticulata* sex chromosome (chromosome 12) and the syntenic *X. maculatus* chromosome 8, particularly in the region of the predicted guppy Stratum I (Fig. 4), which may explain the shifted position of the region with low male coverage in analyses that use the *X. maculatus* genome. In addition, different methodological parameters can have a significant impact on the proportion of reads mapped. Mapping efficiency is substantially reduced when using the *X. maculatus* reference (Table 2), which decreases power to detect a signal of X-Y differentiation.

**Table 2.** Percentage of concordant, properly paired* read alignments.

| Data | Sequencing data | *P. reticulata* |  | *P. wingei* |  |
| --- | --- | --- | --- | --- | --- |
|  | Genome assembly | *P. reticulata* | *X. maculatus* | *P. wingei* | *X. maculatus* |
| Method | bwa -aln/-sampe<br>*Wright et al. 2017* | 59 | 5 | 75 | 17 |
|  | bowtie2 -local<br>*Kirkpatrick et al. 2020* | 83 | 56 | 87 | 71 |
|  | bwa -mem -M<br>*Bergero et al. 2019* | 83 | 72 | 84 | 79 |

\*Both mates of a read pair map to the same chromosome or scaffold, with the expected insert size and read orientation.

**Figure 4.**
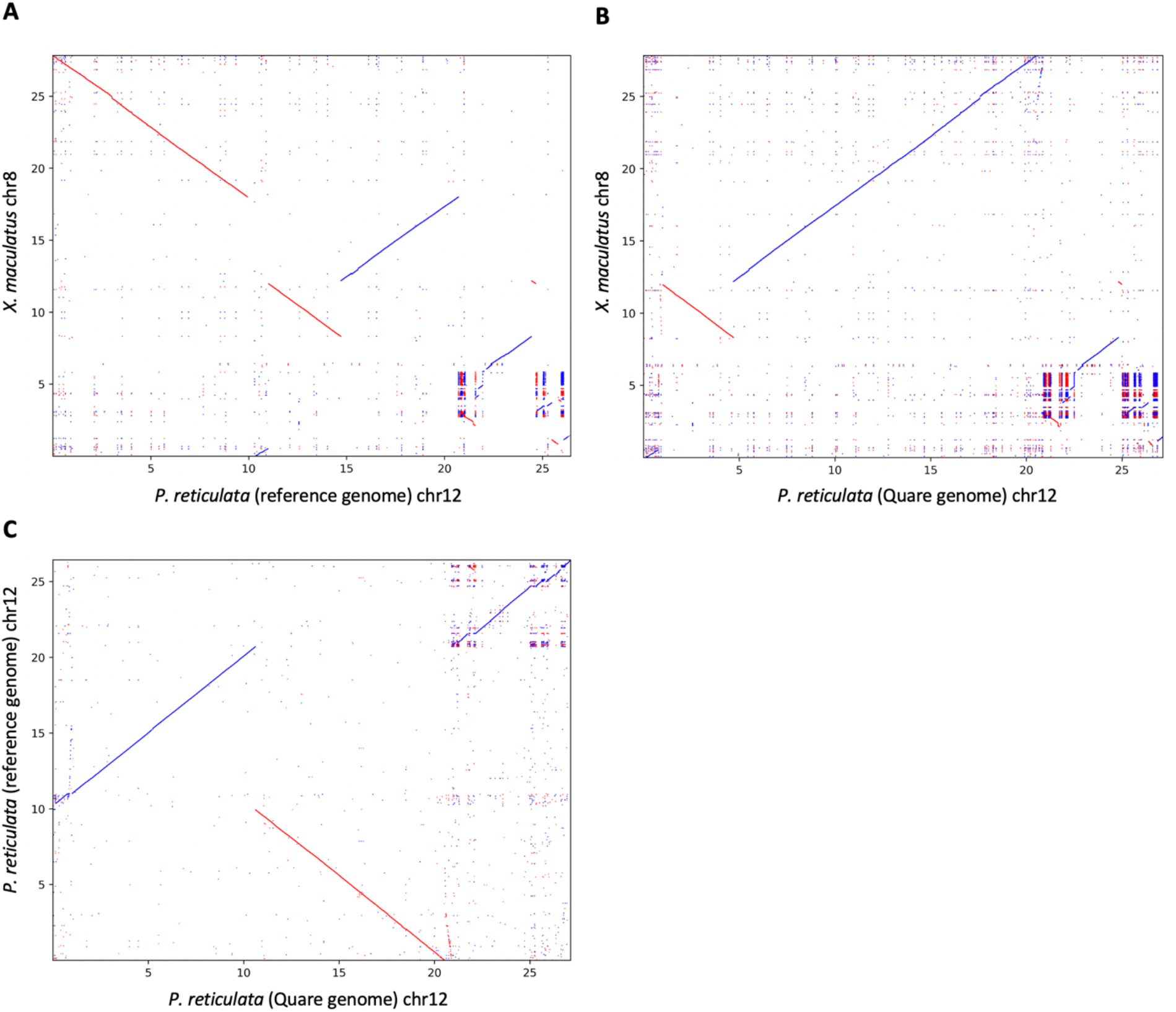
Structural rearrangements and duplications between *P. reticulata* and *X. maculatus* genomes. Dot-plots of alignments between (A) *X. maculatus* chromosome 8 and *P. reticulata* chromosome 12 from the reference genome assembly (Kunstner et al. 2016), (B) *X. maculatus* chromosome 8 and *P. reticulata* chromosome 12 from the Quare *de novo* genome assembly (Almeida et al. 2021), and (C) *P. reticulata* chromosome 12 from the reference genome assembly and chromosome 12 from the Quare *de novo* genome assembly. Forward alignments are shown in blue and reverse alignments in red.

We find no clear pattern of Stratum I when mapping reads to the *P. reticulata* reference genome based on the methodology in Bergero et al. (2019) (Fig. 2C). While we cannot disentangle between the reference genome used and the methodology in this analysis, our other data suggests that the absence of a Stratum I signal is largely due to the *P. reticulata* reference genome. Specifically, when mapping *P. reticulata* reads to the *X. maculatus* genome we recover qualitatively the same coverage pattern across all three methodological approaches (Fig. 2D, E, F). Similarly, our *P. wingei* analyses detailed below reveal that, when using the same genome, the Bergero et al. (2019) pipeline produces very similar patterns to the other two methodological approaches (Fig. 3). Previous work has reported several inversions and assembly errors on the sex chromosome of the first draft of the *P. reticulata* reference genome (Bergero et al. 2019; Darolti et al. 2020; Charlesworth et al. 2020; Fraser et al. 2020), which may be obscuring a signal of Stratum I.

The analyses for *P. wingei* also reveal a lower male to female coverage at the distal end of the chromosome, however, this pattern is only observed in analyses that mapped reads to the *de novo P. wingei* assembly (Fig. 3A, B, C). By contrast, the analyses that used the *X. maculatus* genome all show a significantly elevated read depth in males compared to females, similar to the results in Kirkpatrick et al. (2020) (Fig. 3D, E, F). Previous cytogenetic work has shown that the *P. wingei* Y chromosome is the largest chromosome in the genome, having accumulated a large heterochromatin block (Nanda et al. 2014). However, in addition to the expansion of repetitive sequence, duplication events from the rest of the genome could have also contributed to the remarkable size of the *P. wingei* Y chromosome. Duplications from the X chromosome to the Y chromosome would explain a signal of elevated coverage in males relative to females in this species.

Regions of the sex chromosomes where recombination has recently been halted or greatly suppressed still retain a high degree of similarity between X and Y sequences. They are also expected to show an elevated SNP density in males compared to females, as Y-linked reads carrying Y-specific polymorphisms will still align to the homologous X region of the female reference genome (Fig. 1; Vicoso et al. 2013). We observed this pattern in *P. wingei* (Darolti et al. 2019) and in replicate upstream populations of *P. reticulata* (Wright et al. 2017; Almeida et al. 2020), and we designated this as Stratum II. It is important to note that in contrast to Stratum I, Stratum II appears to have formed independently several times. Therefore, to further quantify divergence between the sex chromosomes we investigated SNP density differences between the sexes using several methodological approaches. In *P. reticulata*, we observe a significantly elevated male SNP density on the sex chromosomes in both of the analyses that aligned reads to the *de novo P. reticulata* genome (Wilcoxon rank sum test *p* < 0.001, Fig. 5A, B). By contrast, the SNP density profiles of the autosomes and the sex chromosomes were indistinguishable in all the analyses that used *X. maculatus* as the reference genome (Fig. 5D, E, F), due to the accumulation of numerous fixed differences between *P. reticulata* and *X. maculatus* which conceal the subtle polymorphisms differences between *P. reticulata* males and females. The *P. wingei* X and Y chromosomes have previously been suggested to be more diverged than those of *P. reticulata*, as shown through more pronounced coverage and SNP density differences between the sexes (Darolti et al. 2019) and a greater accumulation of repetitive sequences on the sex chromosomes in *P. wingei* compared to *P. reticulata* (Morris et al. 2018; Almeida et al. 2021). Our results here confirm this, as we find a significantly higher male:female SNP density for the sex chromosomes compared to the autosomes across all methodological analyses, as well as when using either one of the *P. wingei de novo* or the *X. maculatus* genomes (Wilcoxon rank sum test *p* < 0.001, Fig. 6).

**Figure 5.**
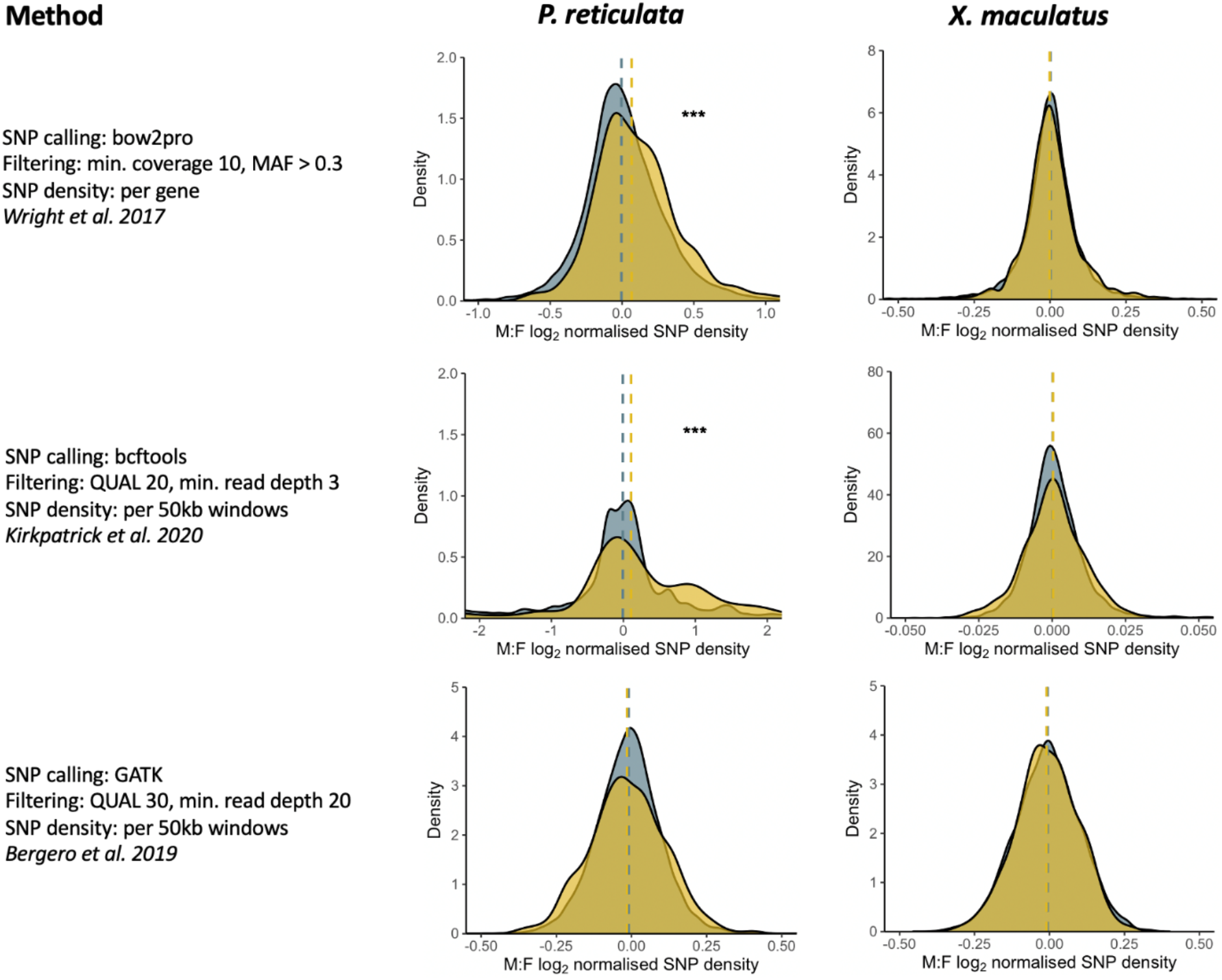
Distribution of *P. reticulata* male:female SNP density for the autosomes (gray) and the sex chromosomes (yellow). Dashed vertical lines indicate median SNP densities and significant differences between the autosomes and the sex chromosomes are shown (*** *p*-value < 0.001).

**Figure 6.**
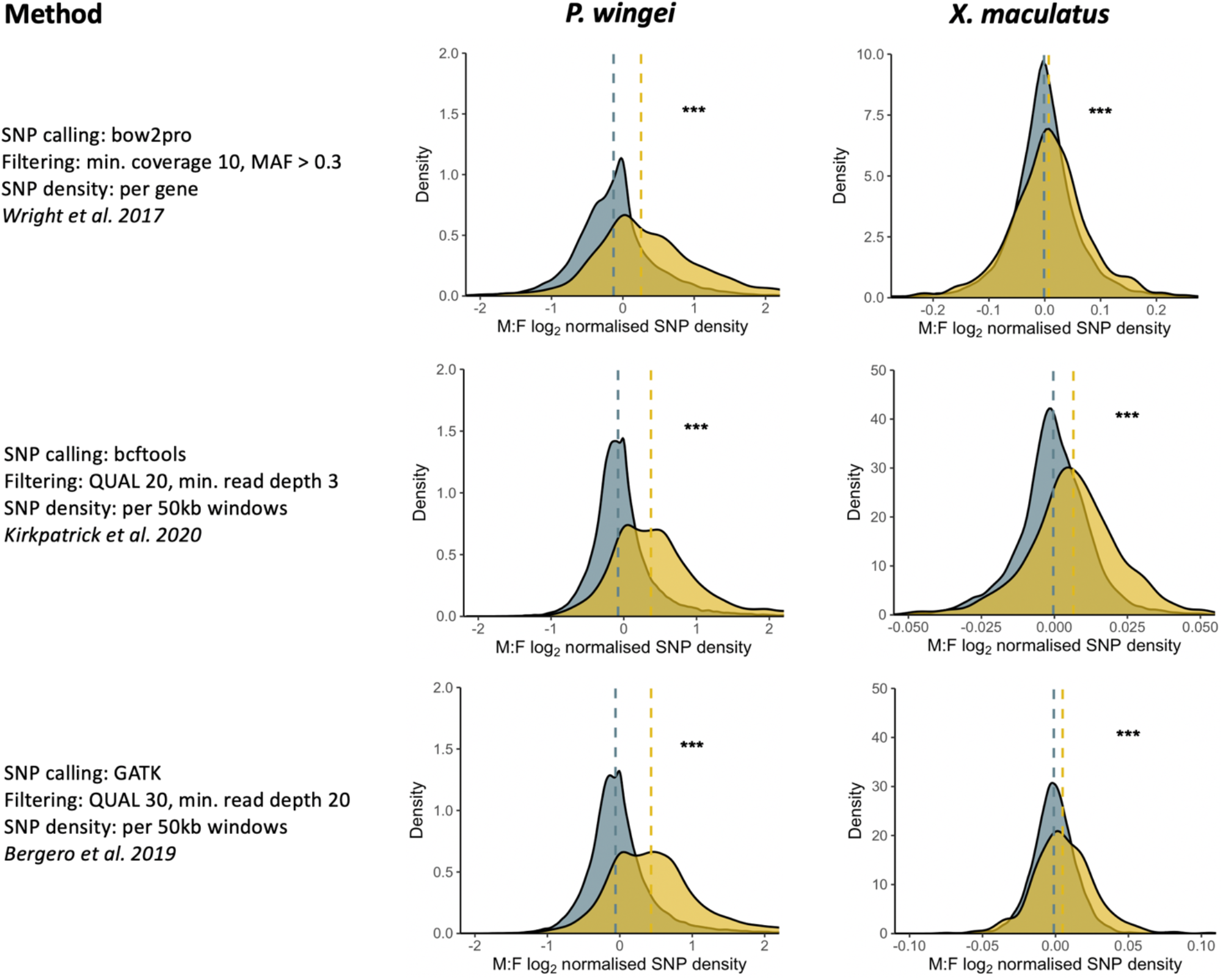
Distribution of *P. wingei* male:female SNP density for the autosomes (gray) and the sex chromosomes (yellow). Dashed vertical lines indicate median SNP densities and significant differences between the autosomes and the sex chromosomes are shown (*** *p*-value < 0.001).

## Discussion

Replication is fundamental to scientific pursuits, and confirmation is necessary to build a robust understanding of the natural world. The expansion of public data efforts has greatly aided transparency and replication efforts, and this remarkable and rapid shift in the scientific culture is exemplified by genomics research, where most of the major journals require deposition of sequencing data as a condition of publication. Failures to replicate results are concerning, and necessitate further work to validate or nullify. However, it is important to understand that different replication approaches will have different risks of Type II errors, or erroneous negative results. This is especially problematic for the detection of subtle, small effect patterns, such as with initial divergence between X and Y chromosomes.

Here we used the same dataset across various methodologies and genome assemblies to test the sensitivity and accuracy of different approaches. Our results show how small changes in the precision of methods can lead to the failure to detect patterns of sex chromosome differentiation in the guppy. The low overall divergence between the X and Y can make detection difficult, but it has nonetheless been observed across multiple datasets, spanning DNA, RNA and methylation data, as well as multiple methods, including comparisons of male and female coverage and SNP density (Wright et al. 2017; Darolti et al. 2019; Almeida et al. 2021), identification of male-specific sequence (Morris et al. 2018; Almeida et al. 2021), phylogenetic analysis of recombination suppression (Darolti et al. 2020; Almeida 2021), and comparative epigenomics (Metzger et al. 2020) (Fig. 7).

**Figure 7.**
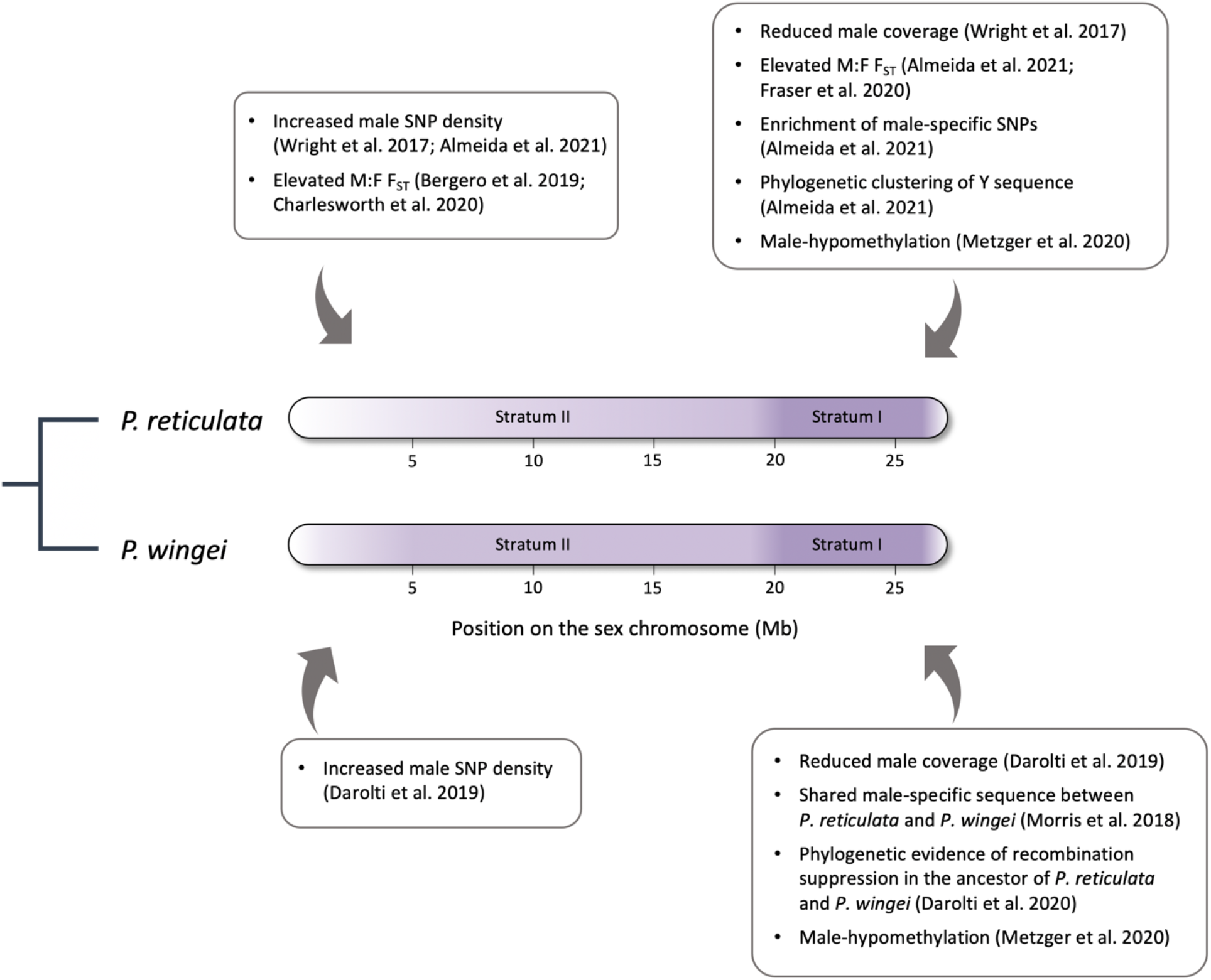
Structure of the *P. reticulata* and *P. wingei* sex chromosomes as predicted by multiple methods, including comparisons of male and female coverage, SNP density, phylogenetic and methylation analyses.

By using the same sample data across multiple methods and genomes, our results illustrate how important methodological differences can alter perceived results, and highlight the need for replication studies to at minimum replicate the analysis using identical methods on the original or equivalent dataset. When possible, replication efforts should go beyond minimum, and expand the analysis by employing more sophisticated methods on existing or expanded datasets. Despite this, some replication efforts use less sophisticated approaches, and in these cases, there is a real concern that a perceived failure of replication is instead the result of a lack of precision or statistical power. This is particularly problematic in the field of genomics, as there is little consensus about the gold standard in methodologies, particularly with regard to data processing and filtering procedures. The lack of standardized practices, coupled with the rich nature of genomic data and the complexity of genomes can make it difficult to discern subtle but important patterns.

Our approach of evaluating the same underlying data with multiple methods and genomes does not account for natural variation across samples and populations, which is substantial (Wright et al. 2017; Almeida et al. 2021). For our *P. reticulata* samples, we chose individuals from an upstream low predation Quare population which we have previously shown to have an intermediate signal of sex chromosome divergence (Wright et al. 2017; Almeida et al. 2021). Samples from populations with greater or lesser signal, or sampling variation due to differences in inversions, duplications and divergence among individuals may also contribute to observed differences.

### Stratum I

We have previously observed evidence for a small region of ancestral recombination suppression in *P. wingei* and *P. reticulata* (Wright et al. 2017; Darolti et al. 2019; Almeida et al. 2021). This has been replicated in some studies, for example Fraser et al. (2020) also found evidence of small regions of Y divergence, and Kirkpatrick et al. (2020) confirmed the phylogenetic clustering of Y sequence in this stratum. However, other studies (Bergero et al. 2019; Charlesworth et al. 2020; Kirkpatrick et al. 2021) did not fully replicate these findings.

It is worth noting that Stratum I region of the guppy Y chromosome is enriched for repetitive elements (Almeida et al. 2021), and reads from this region may, depending on the parameters used, map to repetitive elements across the genome, obscuring real read depth differences between males and females if non-coding sequence is included in the analysis. Focusing on uniquely mapping reads when comparing coverage differences between males and females can minimize issues associated with Y repetitive regions. However, our comparative analysis revealed that a pattern of X-Y differentiation can still be recovered without restricting the analysis to uniquely mapping reads (Fig. 2, Fig. 3). More stringent SNP filtering parameters can also help eliminate noise in genomic comparisons, and this is particularly important when studying young sex chromosomes as they are expected to exhibit subtle divergence signatures. We were, however, able to identify a signal of elevated male SNP density on the sex chromosomes relative to the autosomes, indicative of Y divergence, across several methodological approaches using different degrees of filtering stringency (Fig. 5, Fig. 6).

Beyond mapping parameters, by far the most substantial source of variation in the results of the different pipelines we compared lies in the reference genome used. This is in part due to the extensive structural variation across populations and species (Fig. 4), but also due to sequence evolution. These two factors combined mean that error compounds over phylogenetic distances, and as the distance between the samples and the genome they are mapped to increases, the ability to detect reduced male:female read depth decreases. This is most evidenced in the strategy by Kirkpatrick et al. (2020), who mapped reads from *Poecilia* species to the *Xiphophorus* genome. They argued that changes over the 40 my phylogenetic distance separating these genera was outweighed by the fact that the *Xiphorphorus* genome is more complete. However, the read mapping rate in Table 2 reveals instead that this strategy is less accurate than using less-complete species- or population-specific genome assemblies, as a significantly smaller proportion of *Poecilia* reads map to the *Xiphophorus* genome across all methods, thereby reducing usable data. This problem is exacerbated by the substantial structural differences between *Xiphophorus* and *Poecilia* on the sex chromosome (Fig. 4), further complicating the comparison. Interestingly, their mapping and filtering methods would have detected Stratum I if they had mapped to a con-specific genome (Fig. 2B, Fig. 3B). To a lesser extent, this is also a problem when mapping data to genomes assembled on different *P. reticulata* populations. The genome used can also greatly affect the perceived patterns of SNP diversity, and relying on the distantly related *Xiphophorus* genome can obscure a signal of elevated male SNP density on the sex chromosomes due to fixed differences between the target species reads and the *Xiphophorus* sequence (Fig. 5).

### Stratum II

As recombination is increasingly suppressed in nascent regions of a sex chromosome, we expect the accumulation of Y-specific SNPs, and we observed this in replicate upstream populations of *P. reticulata* (Wright et al 2019; Almeida et al. 2021) and in *P. wingei* (Darolti et al. 2019), consistent with convergent evolution across populations and species (Darolti et al. 2020). Whether this is due to the important environmental effects on recombination rate (Plough 1917; Grell 1971; Stevison et al. 2019), sexual conflict (Wright et al. 2017), neutral shifts in male recombination hotspots (Wright et al 2016; Bergero et al. 2019) or selection against recombinants in the wild remains an important area of further work.

Additionally, given that many mechanisms of recombination suppression only accumulate over time (Furman et al. 2020 and references cited), it also remains unclear how complete recombination suppression is in this region, and whether rare recombination events observed in this region in lab-reared males (Bergero et al. 2019) occur in wild populations. Regardless, it is important to note that suppressed recombination does not necessarily mean that recombination never occurs between the X and Y chromosomes, but rather that it is at least exceedingly rare or recombinant individuals are selected against.

Because of the expected heterogeneity observed in the initial stages of the divergence process (Bergero et al. 2013; Natri et al. 2013; Reichwald et al 2015), sliding window approaches may be insufficient to reveal overall patterns of elevated male SNP density expected in these regions. Density distributions or direct statistical comparisons between species may be required. This is evidenced by our observation of elevated male:female SNP density across nearly all methods (Fig. 5 and 6), with the exception of *P. reticulata* data mapped to the *Xiphophorus* genome, again illustrating the problems with mapping over vast evolutionary distances.

### Concluding remarks

Here we have used the same data to compare methods and genomes in the discovery of nascent sex chromosomes. We hope that our results provide a gold standard for future work in other study systems, and resolve some of the recent controversy over the sex chromosomes in *Poecilia*.

## Acknowledgements

We thank Jacelyn Shu of jacelyndesigns.com for drafting Fig. 1. We gratefully acknowledge funding from the ERC (grant agreement 680951), NSERC and a Canada 150 Research Chair to JEM.

